# The role of cell growth rate on accumulation of the mitotic cyclin Cdc13 in fission yeast

**DOI:** 10.64898/2026.05.14.724355

**Authors:** Sarah E. Vandal, Sayeh Rezaee, Cesar Nieto, Mackenzie J. Flynn, Abhyudai Singh, James B. Moseley

**Author notes:** Address correspondence to: James B. Moseley.

## Abstract

Eukaryotic cells control their size by coordinating growth and division. Fission yeast divide at a reproducible cell size due to regulated activation of the cyclin-dependent kinase Cdk1. The nuclear concentration of mitotic cyclin Cdc13 increases in a time-dependent manner to promote Cdk1 activation as cells grow. Here, we show that interphase Cdc13 is stable against degradation and nuclear export, but is diluted by cell growth. Low glucose reduced cell growth rate but not time-dependent accumulation of Cdc13. Uncoupling the rates of cell growth and Cdc13 accumulation resulted in higher concentrations of nuclear Cdc13 despite reduced cell size. This change coincided with reduced activating phosphorylation of Cdk1-T167. Mathematical modeling and experiments showed that cells maintain size homeostasis under these conditions. In contrast to low glucose, poor nitrogen reduced both cell growth rate and Cdc13 accumulation rate. Therefore, Cdc13 accumulation is independent of cell growth rate but can be altered by nutrient-specific mechanisms.

## Introduction

The size of a dividing cell typically scales with its growth rate. From bacteria to human cells, reduction in growth rate results in smaller cell size (Schaechter et al., 1958; Fantes and Nurse, 1977; Johnston et al., 1979; Pierucci, 1978; Kellogg and Levin, 2022). This correlation between cell growth rate and cell size suggests a mechanistic link. Some examples have been identified including bacterial cell division proteins DnaA and FtsZ and budding yeast cyclin protein Cln3, whose expression correlates with cell growth rate under multiple nutrient-limited conditions (Sommer et al., 2021; Schneider et al., 2004; Hall et al., 1998; Gallego et al., 1997; Weart and Levin, 2003). However, many aspects of the link between cell growth rate and cell size remain incompletely understood.

The fission yeast *Schizosaccharomyces pombe* serves as a strong model system to study connections between cell size and growth rate. Like other organisms, the growth rate of fission yeast cells can be modulated by nutrient limitation. In nutrient rich conditions, fission yeast cells exhibit strong size homeostasis in which all cells grow at the same rate and divide at the same size, regardless of their size at birth (Fantes, 1977). In nutrient poor conditions, fission yeast cells grow at a reduced rate and divide at a smaller size. This phenomenon has been observed in multiple conditions including limitations to carbon (e.g. reduced glucose) and nitrogen (Fantes and Nurse, 1977; Saitoh and Yanagida, 2014; Bertaux et al., 2023). Both glucose and nitrogen limitation reduce cell growth rate and cell size, but these different nutrient conditions act primarily through different signaling pathways (Hardie and Carling, 1997; Uritani et al., 2006; Davie et al., 2015; Ling et al., 2020). How nutrient limitation connects with cell size and cell growth rate remains an open mechanistic question.

The size of fission yeast cells depends primarily on a cell size checkpoint that operates at the G2/M transition, which is controlled by the ubiquitous cyclin-dependent kinase Cdk1 (Nurse and Thuriaux, 1980). Cdk1 activity is partly controlled through inhibitory phosphorylation of Cdk1-Y15 and activating phosphorylation of Cdk1-T167 (Gould et al., 1991; Hermand et al., 2001). In particular, Cdk1-pY15 contributes to cell size control and is regulated by the kinase Wee1 and phosphatase Cdc25 (Russell and Nurse, 1987, 1986; Gould and Nurse, 1989; Moreno et al., 1990; Gautier et al., 1991). The cellular activities of Wee1 and Cdc25 scale with cell size to promote size-dependent activation of Cdk1 (Pan et al., 2014; Miller et al., 2023; Facchetti et al., 2019). Nutrient sensing pathways regulate Wee1 and Cdc25, providing an initial molecular connection between nutrients, cell size, and Cdk1 activation (Yanagida et al., 2011; Allard et al., 2019; Belotti et al., 2011).

Activation of Cdk1 also requires binding to a cyclin subunit, and the B-type cyclin Cdc13 is the only fission yeast cyclin that is both necessary and sufficient to drive the mitotic cell cycle through Cdk1 (Fisher and Nurse, 1996, 1995; Coudreuse and Nurse, 2010). The nuclear concentration of Cdc13 increases during the cell cycle to promote increased activation of Cdk1 in the nucleus, eventually leading to initiation of the G2/M transition (Miller et al., 2023; Curran et al., 2022; Sugiyama et al., 2024; Bashir et al., 2026 Preprint). Recent studies have shown that Cdc13 accumulation scales with time and not cell size (Miller et al., 2023; Sugiyama et al., 2024), although effects from cell size have also been reported (Bashir et al., 2026 Preprint).

In this study, we defined the mechanisms of time-dependent accumulation of Cdc13 in the nucleus, and then tested if this accumulation rate depends on cell growth rate. We found that glucose limitation reduces the rate of cell growth but not the time-dependent rate of Cdc13 accumulation, leading to high nuclear concentration of Cdc13 in low glucose conditions. This time-dependent accumulation occurs through constant accumulation of Cdc13, which is stable against degradation and nuclear export during interphase. Unlike glucose limitation, poor nitrogen reduced the rate of Cdc13 accumulation. Thus, the rate of Cdc13 accumulation does not depend strictly on cell growth rate but can instead be regulated by nutrient-specific signals. Finally, we integrated these dynamic, single-cell measurements into a mathematical model. We demonstrate that the observed changes in cell size regulation can be explained by altering just two parameters in low glucose: a reduced cell growth rate and a shift in the Cdk1 activation threshold.

## Results and Discussion

### Cdc13 protein is stable in the nucleus

Cdc13 accumulation can be regulated by changes in Cdc13 synthesis, degradation, nuclear transport, or dilution by cell volume expansion. To establish the baseline parameters of Cdc13 accumulation, we first examined the contribution of *cdc13* transcription using single-molecule FISH. Consistent with other work (Bashir et al., 2026 Preprint), we found that the concentration of the *cdc13+* transcript is constant across cell size (Fig. 1A-B), indicating that Cdc13 protein accumulation is controlled post-transcriptionally. Next, we investigated the half-life of Cdc13 protein to test the role of Cdc13 protein degradation. We used single-cell timelapse microscopy to measure the nuclear concentration of an internally tagged and functional version of Cdc13-sfGFP expressed at the endogenous *cdc13^+^* locus (Miller et al., 2023; Kamenz et al., 2015). We measured Cdc13 degradation following the addition of the protein synthesis inhibitor cycloheximide (CHX) (Fig. 1C-E, S1A). We found that Cdc13 is a stable protein in the nucleus during interphase, with a half-life of approximately 173 minutes (Fig. 1E).

**Figure 1.**
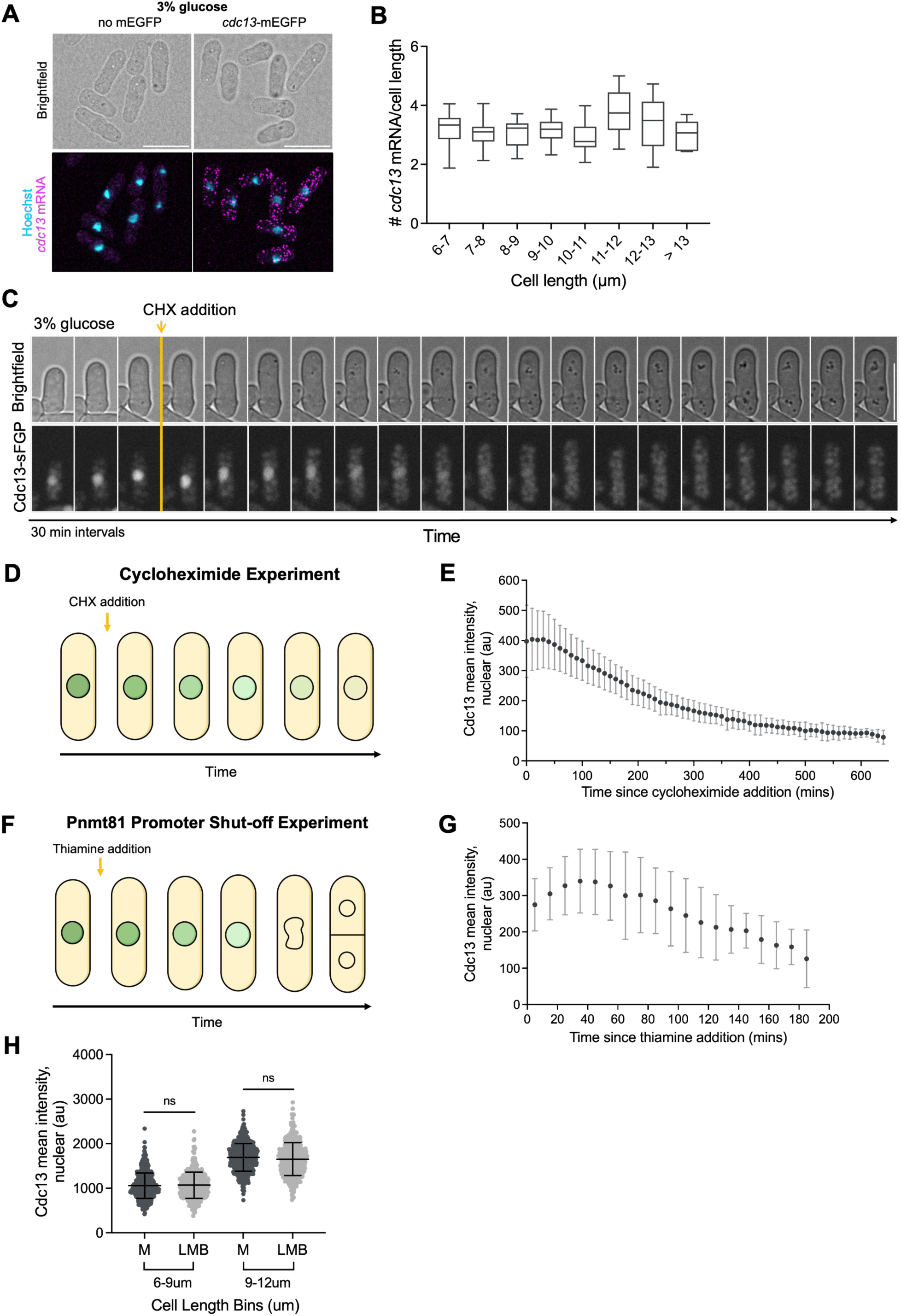
Cdc13 protein is stable in the nucleus. (A) Representative brightfield and smFISH *cdc13-mEGFP* transcript fluorescence images of *cdc13-mEGFP-3’UTR (cdc13)* cells grown in 3% glucose. Hoechst stain marks the nucleus (scale bar = 10 µm). (B*) cdc13+* mRNA concentration (# molecules per cell length) plotted as a function of cell size in 1 µm size bins for cells grown in 3% glucose. (C) Representative brightfield and fluorescence timelapse images of an interphase *cdc13-sfGFP* cell during cycloheximide treatment. (D) Schematic of Cdc13-sfGFP cycloheximide experiment. (E) Nuclear Cdc13-sfGFP concentration (au) of single cells treated with cycloheximide (1000 µg/mL) over time (n = 24 cells). 0 mins indicates cycloheximide addition. Timepoints were fit to a one-phase decay function to determine half-life of Cdc13 (t1/2 = 173 mins). (F) Schematic of *Pnmt81-cdc13-sfGFP* shut-off experiment. These cells enter mitosis due to expression of *cdc13+* from the endogenous locus. (G) Nuclear Cdc13-sfGFP concentration (au) of *Pnmt81-cdc13-sfGFP* single cells exposed to thiamine (2 µg/mL) over time (n = 20 cells). 0 mins indicates thiamine addition. (H) Mean nuclear Cdc13-sfGFP concentration (au) of interphase cells treated with leptomycin B (50 ng/µL) or methanol control after 60 mins grown in 3% glucose media. ns p > 0.05 by unpaired t-test.

We then performed two complementary tests of Cdc13 stability. First, we measured untagged Cdc13 in CHX-treated cells arrested in G2 by the conditional *cdc25-22* mutation and observed a long half-life (Fig. S1B-C). Second, we constructed a strain in which the endogenous *cdc13+* gene was unmodified, but a second copy of Cdc13 including a fluorescent tag was integrated at the *his5+* locus and expressed by the thiamine-repressible *P81nmt1* promoter (Maundrell, 1990; Basi et al., 1993). We used timelapse microscopy to measure the rate of Cdc13-sfGFP degradation after addition of thiamine and before mitotic entry. This experiment yielded a slow rate of Cdc13 degradation (Fig. 1F-G, S1D), similar to our results with CHX. These combined data suggest that time-dependent accumulation of Cdc13 occurs post-transcriptionally through steady synthesis of a stable protein. Lastly, we tested the contribution of nuclear export on Cdc13 accumulation using the nuclear export inhibitor Leptomycin B (LMB). We compared Cdc13 nuclear concentration following treatment with either LMB or methanol (vehicle control) and found that LMB did not affect the nuclear concentration of Cdc13 in either small cells or large cells (Fig. 1H). This suggests that active nuclear export does not counteract the accumulation of Cdc13 during interphase. Together, our results indicate a time-dependent accumulation mechanism of constant synthesis of stable Cdc13 protein in the nucleus.

### Glucose limitation slows cell growth but not Cdc13 accumulation

If the nuclear concentration of Cdc13 is not counteracted by degradation or export during interphase, then dilution through cell growth is the primary mechanism counteracting its time-dependent accumulation. Nuclear size scales with cell size across a range of conditions (Neumann and Nurse, 2007), which means that slowing cell growth rate should slow nuclear growth and therefore Cdc13 dilution. In budding yeast, G1 cyclin Cln3 concentration scales with cell growth rate (Sommer et al., 2021; Schneider et al., 2004; Hall et al., 1998). Therefore, we tested if Cdc13 concentration is similarly growth rate dependent by altering glucose availability. We compared cells grown in standard high 3% glucose (167 mM) or low 0.08% glucose (4.4 mM). As expected, cells in low glucose grew at a slower rate and divided at a smaller size compared to cells grown in high glucose (Fig. 2A-B, S1E) (Schaechter et al., 1958; Fantes and Nurse, 1977; Johnston et al., 1979; Petersen and Nurse, 2007; Pluskal et al., 2011). We used the same internally tagged version of Cdc13-sfGFP to measure Cdc13 concentration across the bulk cell population. Interestingly, Cdc13 nuclear concentration was dependent upon glucose availability (Fig. 2C, S1F). For a given cell size, cells growing slowly in low glucose had a higher nuclear concentration of Cdc13 compared to cells growing rapidly in high glucose (Fig. 2C, S1G-H). These data suggest that the mechanisms controlling Cdc13 accumulation do not scale with cell growth rate under glucose limited conditions.

**Figure 2.**
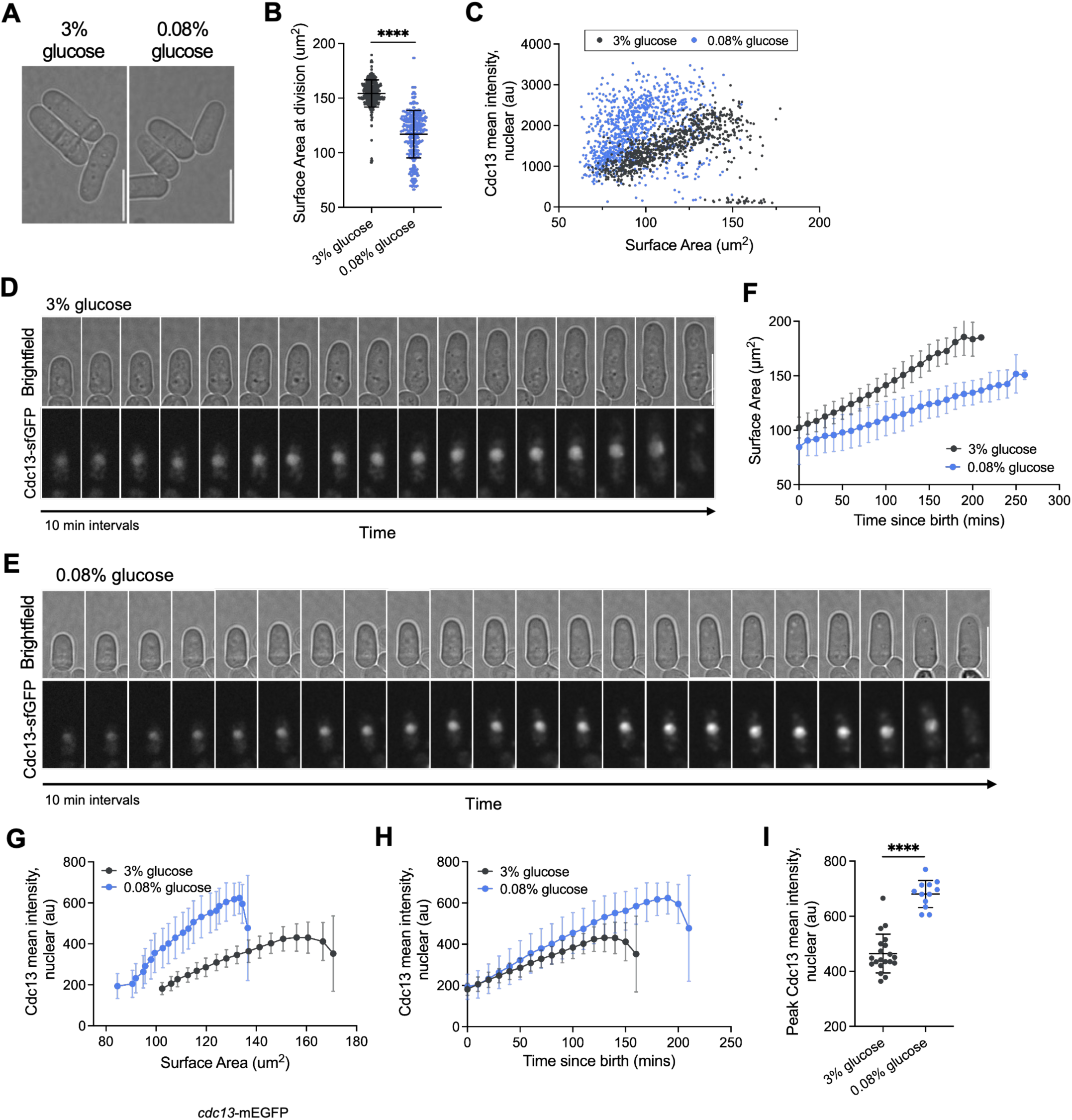
Low glucose reduces cell growth rate but not Cdc13 nuclear accumulation. (A) Brightfield images of wildtype cells grown in rich media with 3% or 0.08% glucose (scale bar = 10 µm). (B) Cell size at division (µm^2^) of *cdc13-sfGFP* cells grown in rich media with 3% or 0.08% glucose (n = 458 cells for 3% glucose; n = 384 cells for 0.08% glucose). (C) Cdc13-sfGFP nuclear concentration (au) for cells plotted by cell size (µm^2^). Cells were grown in 3% glucose (n = 843 cells) or 0.08% glucose (n = 987 cells). (D-E) Representative brightfield and fluorescence timelapse images of a *cdc13-sfGFP* cell grown in 3% glucose (panel D) or 0.08% glucose (panel E). (scale bars = 10 µm). (F) Quantification of cell growth rate for cells grown in 3% or 0.08% glucose. Error bars indicate standard deviation (3% glucose n = 20 cells; 0.08% glucose n = 12 cells). (G-H) Cdc13-sfGFP concentration in the nucleus of single cells from timelapse imaging for cells grown in 3% or 0.08% glucose. Concentrations are plotted against either cell size (µm^2^) (panel G) or time since birth (panel H). Error bars indicate standard deviation. (I) Peak Cdc13-sfGFP mean nuclear concentration (au) prior to division for cells grown in media with 3% or 0.08% glucose. The same cells are plotted in panels F-I. **** p <0.0001 by unpaired t-test.

To directly measure the rate of Cdc13 accumulation we followed single cells growing in high or low glucose using timelapse microscopy coupled with microfluidics. Similar to our bulk assays, cells in low glucose grew more slowly and divided at a smaller size than cells grown in high glucose (Fig. 2D-F). We compared Cdc13 nuclear concentration over either time since cell birth or cell size. We found that Cdc13 nuclear concentration scales differently with cell size under high and low glucose conditions (Fig. 2G), similar to bulk assays. However, the time-dependent accumulation rate of Cdc13 in the nucleus was nearly identical under high and low glucose conditions (Fig. 2H and S1I). Therefore, cells grown in low glucose grow slower but accumulate nuclear Cdc13 at a similar rate to cells grown in high glucose, resulting in a higher peak concentration of nuclear Cdc13 before entering mitosis (Fig. 2I). Consistent with this model, cells grown in an intermediate concentration of glucose (0.1%, 5.5 mM) accumulated Cdc13 at a similar rate resulting in a peak concentration in between high and low glucose conditions (Fig. S1J-K). These results suggest that glucose availability may tune the threshold of Cdc13 needed for mitotic entry without impacting the rate of Cdc13 accumulation. Together these findings suggest that Cdc13 accumulates as a molecular timer that is independent of cell growth rate.

To test if Cdc13 concentration in low glucose depends on time or cell size, we used large *cdr2Δ* mutant cells that uncouple cell cycle timing and cell size. We measured Cdc13 nuclear concentration in wild-type and *cdr2Δ* cells grown in low glucose, similar to our previous work in high glucose (Miller et al., 2023). In these two strains, the concentration of Cdc13 scaled identically as a function of time since birth, despite their different sizes (Fig. S1L). Accordingly, the concentration of Cdc13 scaled differently for wild-type and *cdr2Δ* when plotted against cell size (Fig. S1M). These results confirm that Cdc13 concentration increases in a time-dependent manner for cells grown in high glucose versus low glucose, despite their different cell growth rates. Therefore, time-dependent Cdc13 accumulation is uncoupled from cell growth rate by changes in glucose availability.

Lastly, we tested if reduced cell growth rate in low glucose affects the degradation rate of Cdc13 protein or the concentration of *cdc13+* transcript. We added CHX to cells grown in 0.08% glucose media and measured a Cdc13 half-life (165 minutes) that was similar to cells grown in high glucose (Fig. S1N-O). We also examined *cdc13*+ RNA number using smFISH. We found a decrease in the concentration of *cdc13+* RNA in low glucose versus high glucose conditions (Fig. S1P-Q). This decrease was seen for 2 out of 3 housekeeping transcripts tested as controls (Fig. S1R-T). This trend is consistent with the concentration of most budding yeast transcripts scaling with cell growth rate (Gao et al., 2026), although the cell cycle regulator Whi5 is an exception (Qu et al., 2019). Our data support a model where post-transcriptional mechanisms drive Cdc13 accumulation at a constant rate that is counteracted by growth-mediated dilution. When cell growth is reduced under low glucose conditions, the nuclear concentration of Cdc13 is higher due to reduced dilution.

### Cdk1 activating phosphorylation is reduced under glucose limitation

Low glucose reduces the rate of Cdc13 dilution but not rate of Cdc13 accumulation. Thus, cells of a given size have a higher concentration of nuclear Cdc13 in low glucose than in high glucose. However, cells in low glucose do not enter into mitosis and divide when they reach the “threshold” concentration of Cdc13 observed for cells in high glucose. Instead, the nuclear concentration of Cdc13 prior to mitotic entry is greater in low glucose compared to high glucose (Fig. 2I). This suggests that the threshold concentration of Cdc13 required to activate Cdk1 and promote mitosis is increased in low glucose. Since the nuclear concentration of Cdk1 is unaffected by glucose limitation (Fig. 3A), this raises the possibility that other mechanisms affecting Cdk1-Cdc13 catalytic activity might compensate for this increased nuclear concentration of Cdc13 in low glucose. The Cdk1-Cdc13 complex is both positively and negatively regulated by phosphorylation (Gould et al., 1991; Gould and Nurse, 1989). We first examined the levels of inhibitory Cdk1-pY15 phosphorylation, which is mediated by Wee1 kinase, but did not observe differences between high glucose and low glucose conditions (Fig. 3B-C). Next, we tested the levels of activating Cdk1-pT167 phosphorylation (Gould et al., 1991; Hermand et al., 2001) and measured a ∼25% reduction in Cdk1-pT167 in low glucose compared to high glucose (Fig. 3D-E). This result suggests that the fraction of active Cdk1-Cdc13 complex may be reduced in low glucose. This mechanism would increase the threshold concentration of Cdc13 at mitotic entry in low glucose conditions (Fig. 3F).

**Figure 3.**
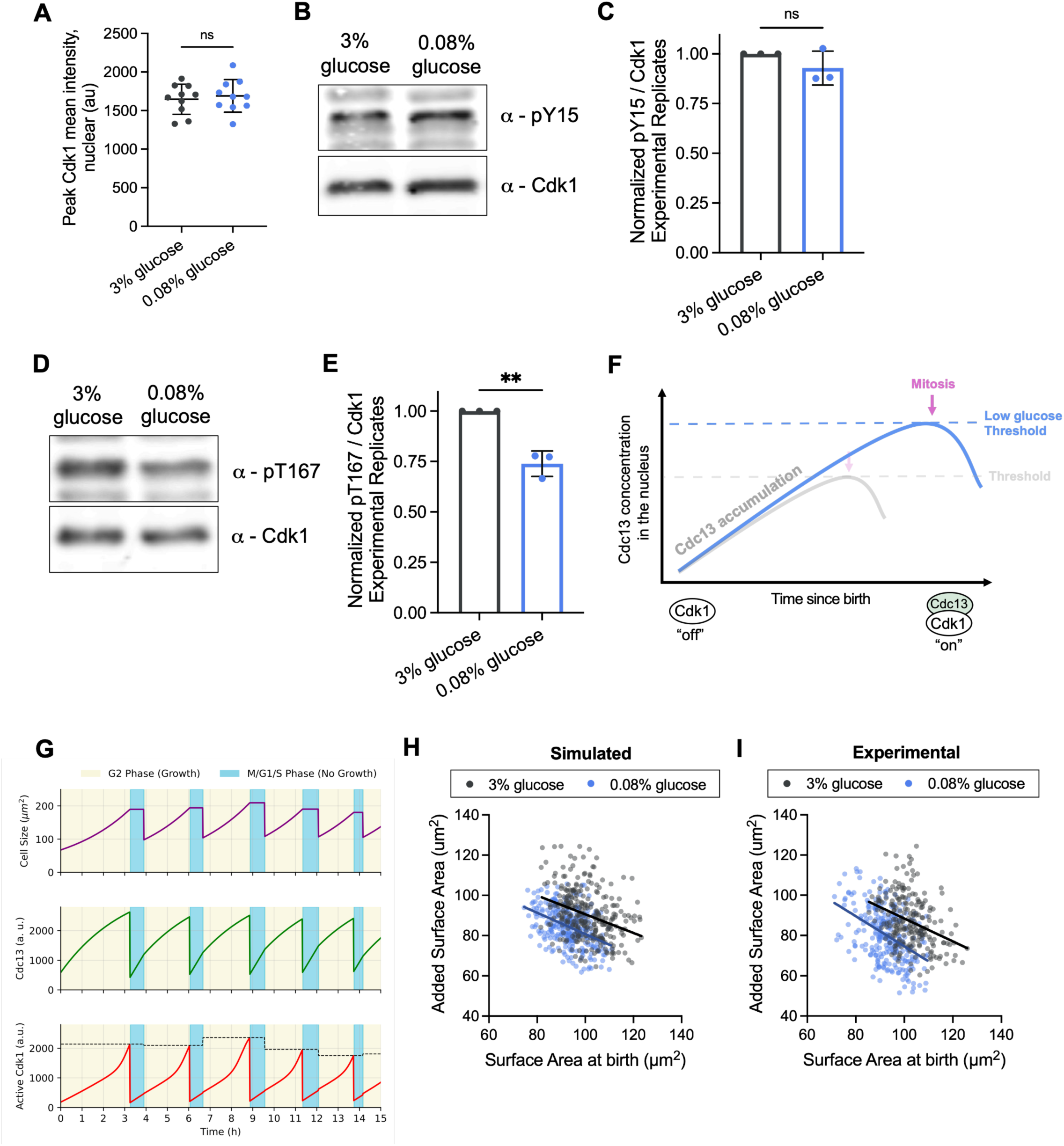
Reduced Cdk1-activating phosphorylation contributes to altered Cdc13 threshold for mitotic entry. (A) Peak Cdk1 mean nuclear concentration (au) prior to division in cells grown in media with 3% or 0.08% glucose (n = 7 cells for each condition). (B) Immunoblot for phosphorylation of Cdk1-Y15 (pY15) for cells grown in 3% or 0.08% glucose. (C) Quantification of Cdk1-pY15 from 3 biological replicates. (D) Immunoblot for phosphorylation of Cdk1-pT167 (pT167) for cells grown in 3% or 0.08% glucose. (E) Quantification of Cdk1-pT167 from 3 biological replicates. ns p > 0.05; ** p < 0.01 by unpaired t-test. (F) Schematic of Cdc13 time-dependent accumulation under high and low glucose. (G) Simulated trajectories of model variables of cell size, Cdc13 dynamics, and active Cdk1 over time. Dashed line in active Cdk1 trajectory indicates the random Cdk1 activation threshold, *c*^†^. Parameters for these example trajectories were taken from the high-glucose condition in Table 1. (H) Simulated size homeostasis plot of cells grown in rich media with 3% or 0.08% glucose. All simulation parameters for different glucose concentrations were kept the same except growth rates and mean Cdk1 activation thresholds (see Table 1). n = 300 simulated cell cycles for each condition. Slopes of linear regression lines are −0.45 for 3% glucose, and −0.53 for 0.08% glucose. (I) Size homeostasis plot of cells grown in 3% or 0.08% glucose (n = 225 cells for 3% glucose; n = 265 cells for 0.08% glucose). Slope of linear regression lines are −0.57 for 3% glucose, and −0.75 for 0.08% glucose.

**Table 1.**
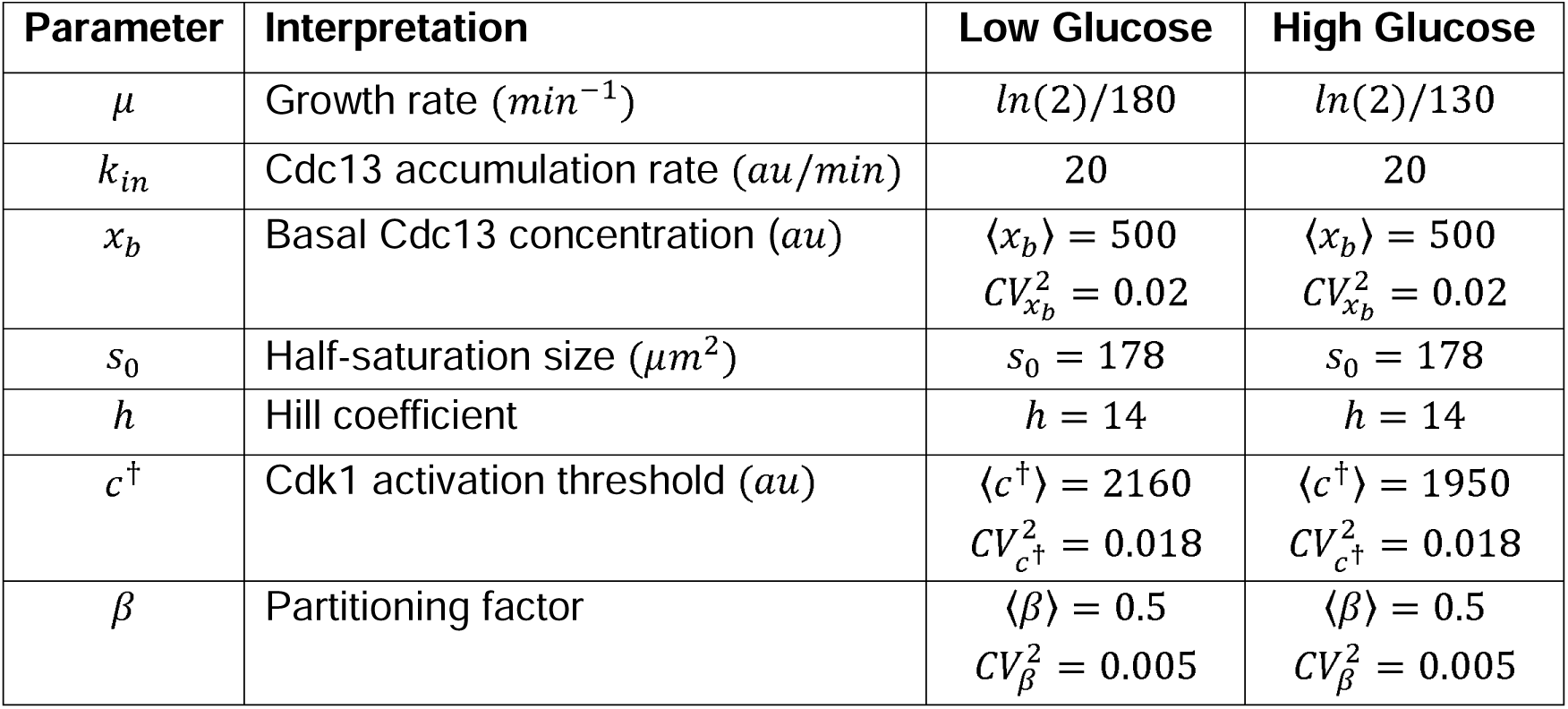
Parameters of the mathematical framework used for modeling division under different glucose concentrations. Parameters 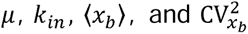 were estimated from experimental data (**Fig. 2**). The remaining parameters were determined by MMD-based optimization.

### Mathematical model of Cdc13 accumulation in high and low glucose

To understand how glucose availability influences Cdc13 dynamics and cell size homeostasis, we generated a minimal mathematical model of the fission yeast cell cycle based on experimental results and previously published work (Miller et al., 2023). Our model describes cell cycle progression through two phases based on measurements of *S. pombe* cell growth (Knapp et al., 2019; Pickering et al., 2019). The first phase is a growth phase (G2), where cells increase exponentially in size over time and Cdc13 accumulates in the nucleus at a constant, size-independent rate with dilution by cell growth. The second phase is a division phase (M/G1/S), where cells stop their growth and Cdc13 levels are reset to a basal value mimicking APC-mediated degradation. Entry into mitosis via Cdk1 activation is dependent on both Cdc13 concentration and size-dependent regulatory inputs mediated by the Cdc25–Wee1 pathway. Random variability in Cdk1 activation threshold and M/G1/S cycle duration timing is included to capture the stochastic nature of the underlying biological processes (Fig. 3G). In our model, the difference between high and low glucose conditions are simplified to two experimentally derived parameters: cell growth rate and the threshold of active Cdk1 required to trigger division. Modeling a nutrient-dependent threshold of active Cdk1 reflects our experimental finding of a reduction in the activating Cdk1-pT167 phosphorylation in low glucose. All other parameters of the model are kept constant across nutrient conditions (Table 1).

We used the model to make predictions about cell size homeostasis in low glucose versus high glucose. Cell size homeostasis can be measured by performing timelapse microscopy on a population of cells, and then plotting each cell’s added cell size versus cell size at birth (“size homeostasis slope”). Fission yeast cells in high glucose have previously been shown to generate a negative slope in these experiments, indicative of strong cell size homeostasis (Sveiczer et al., 1996; Facchetti et al., 2017). Our model predicts that cells in low glucose will retain strong cell size homeostasis despite reduced cell growth rate and increased Cdc13 concentration (Fig. 3H). We tested this prediction experimentally and observed a negative size homeostasis slope under both low glucose and high glucose conditions (Fig. 3I). We conclude that cells maintain strong cell size homeostasis in low glucose despite increased concentrations of Cdc13.

### Cdc13 nuclear accumulation is reduced under nitrogen limitation

Our data show that Cdc13 accumulation and cell growth are uncoupled during glucose limitation. To test if Cdc13 accumulation was similarly uncoupled from growth rate in response to other nutrient limitations, we compared cells grown in a rich nitrogen source (glutamate) versus a poor nitrogen source (proline). Cells in proline grew at a reduced rate and divided at a smaller size than cells in glutamate (Fig. S2A-B) (Petersen and Nurse, 2007; Fantes and Nurse, 1977). We found that Cdc13 accumulated at a slower rate for cells grown in proline compared to those grown in glutamate, when plotted either by time since birth or cell size (Fig. 4A-E). Interestingly, unlike what we observed during glucose limitation, the peak nuclear concentration of Cdc13 prior to cell division was lower in proline versus glutamate (Fig. 4F). We also tested cells grown in isoleucine, which is a worse nitrogen source than proline, and observed an even further decreased peak Cdc13 concentration and cell size (Fig. S2D). These data demonstrate that poor nitrogen reduces both cell growth rate and Cdc13 accumulation rate, and suggest that Cdc13 accumulation is differentially controlled across unique nutrient conditions.

**Figure 4.**
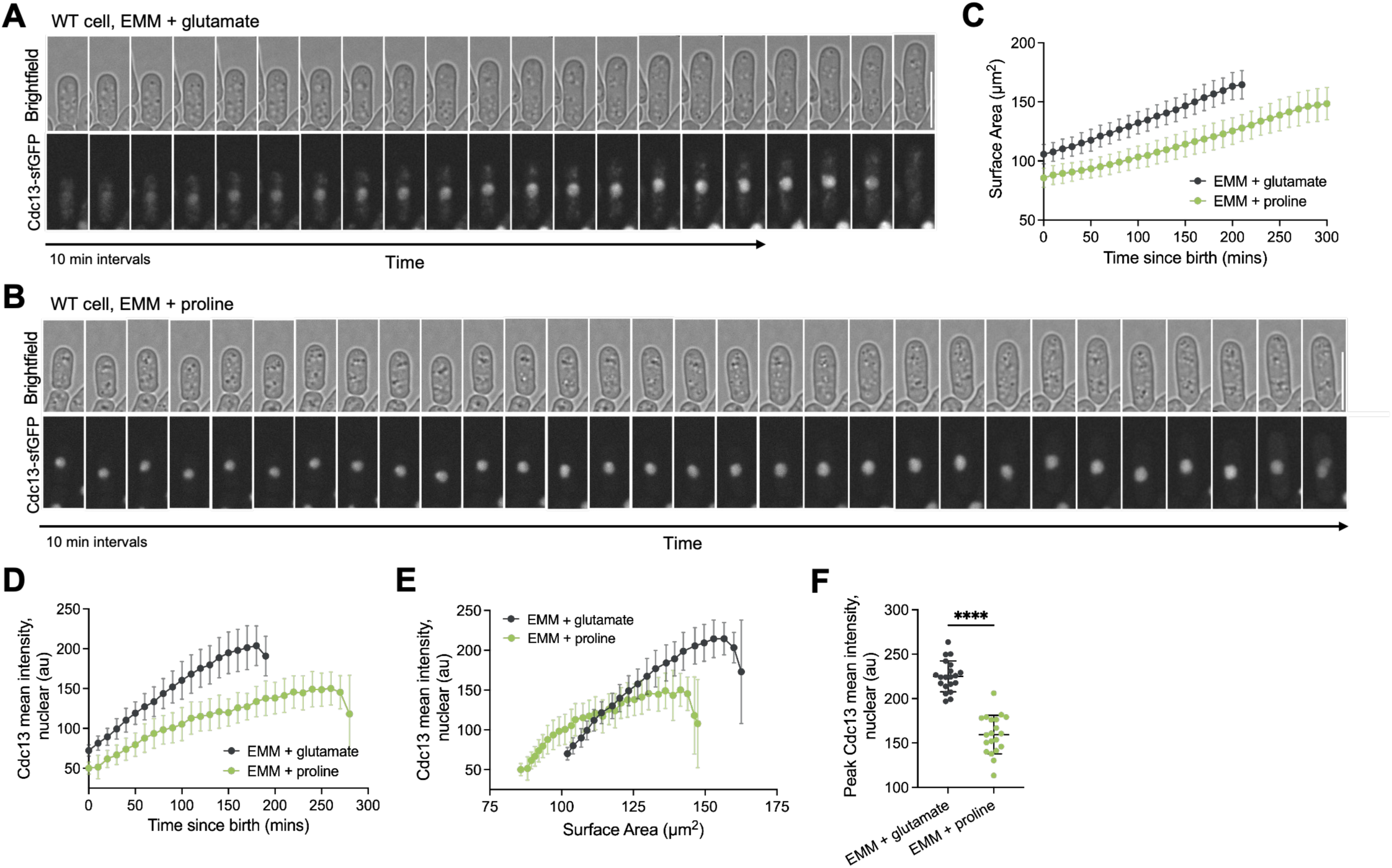
Nitrogen limitation reduces cell growth rate and Cdc13 accumulation rate. (A-B) Representative brightfield and fluorescence timelapse images of a *cdc13-sfGFP* cell grown in minimal media supplemented with either glutamate (panel A, rich nitrogen) or proline (panel B, poor nitrogen) (scale bar = 10 µm). (C) Quantification of cell growth rate for cells grown in minimal media supplemented with glutamate and proline (n = 20 cells for each condition). Error bars indicate standard deviation. (D-E) Cdc13-sfGFP nuclear concentration plotted as a function of either time since cell birth (panel D) or cell size (panel E). Cells were grown in media containing either glutamate or proline as indicated. Error bars indicate standard deviation. (F) Peak Cdc13-sfGFP mean nuclear concentration (au) prior to division for cells grown in media supplemented with glutamate or proline. Same cells are plotted in C-F. **** p < 0.0001 by unpaired t-test.

Finally, we tested if Cdc13 accumulation functions as a strict timer or sizer in poor nitrogen by comparing wild-type cells and large *cdr2Δ* cells. In proline-containing media, wild-type and *cdr2Δ* mutants had different rates of Cdc13 accumulation when plotted by either time or size (Fig. S2E-F), although they were more closely related by time. This result contrasts both high and low glucose, where wild-type and *cdr2Δ* mutants accumulate nuclear Cdc13 at the same time-dependent rate (Miller et al., 2023). Overall, our experiments in nitrogen limitation demonstrate the existence of nitrogen-sensing pathways that alter the rate of nuclear Cdc13 accumulation. More broadly, our results demonstrate that Cdc13 nuclear accumulation can be tuned differently under varied nutrient stresses.

## Conclusions

Our work provides new insights into time-dependent accumulation of the mitotic cyclin Cdc13 under different cell growth conditions. We propose that Cdc13 accumulates during interphase through steady synthesis and nuclear import countered by dilution through cell growth. Activation of the APC during mitosis leads to Cdc13 degradation (Yamano et al., 1996), which resets the system for the subsequent cell cycle. A major conclusion of our work is that Cdc13 accumulation is not strictly tied to cell growth rate. We found that glucose limitation reduced cell growth rate but not Cdc13 accumulation rate. This means that Cdc13 is diluted more slowly without compensatory changes in time-dependent accumulation, leading to a higher Cdc13 concentration for cells grown in low glucose. The separation of cell growth rate from Cdc13 accumulation rate differs from *S. cerevisiae* G1 cyclin Cln3, which appears to be strictly tied to cell growth rate (Sommer et al., 2021; Schneider et al., 2004; Hall et al., 1998; Gallego et al., 1997). This difference raises questions about mechanisms that are specific to different cyclin proteins and organisms. Among *S. pombe* cyclins, Cdc13 is unique in its ability to drive complete mitotic cell cycles in the absence of other cyclins (Fisher and Nurse, 1996). However, other fission yeast cyclins including Cig1, Cig2, and Puc1 are also expressed during mitotic cell cycles. Future studies on various cyclin proteins from different organisms might reveal common principles among the cyclin regulatory networks.

We found that cells grown in low glucose enter mitosis at a higher threshold level of Cdc13, as compared to cells grown in high glucose. Two possibilities that are not mutually exclusive might account for this higher threshold. First, other regulatory mechanisms could reduce the kinase activity of Cdk1-Cdc13. We observed reduced levels of activating T167 phosphorylation in low glucose, consistent with Cdk1-activating phosphorylation scaling with cell growth rate in budding yeast (Blank et al., 2026 Preprint). More broadly, we note that Cdc13 accumulation functions as part of a larger cell size control network that regulates Cdk1 activity through multiple pathways and inputs. Second, the protein phosphatases that counteract Cdk1 might be altered in low glucose. These phosphatases set an inhibitory threshold that must be overcome by Cdk1 activation through multiple mechanisms including Cdc13 accumulation. Nutrient-sensing pathways such as Greatwall-Endosulfine can modify phosphatase activity and represent targets for future studies on this system (Chica et al., 2016; Pérez-Hidalgo and Moreno, 2017).

Our results in nitrogen limitation indicate that Cdc13 accumulation can be altered in a nutrient-specific manner. While glucose limitation uncoupled cell growth rate and Cdc13 accumulation rate, we found that poor nitrogen reduced both cell growth rate and Cdc13 accumulation rate. These results indicate that nutrient-specific signals regulate Cdc13 when glucose versus nitrogen is limited. It is possible that nitrogen-sensing pathways such as TOR reduce Cdc13 accumulation along with cell growth rate. Alternatively, glucose-modulated pathways such as AMPK or PKA could buffer Cdc13 accumulation from reduced cell growth rate. Future studies will be needed to test these and other candidate regulatory systems.

## Materials and Methods

### Yeast strains and growth

Standard *Schizosaccharomyces pombe* media and methods were used (Moreno et al., 1991). Strains used in this study are listed in Supplemental Table 1. Gene tagging and deletion were performed using PCR and homologous recombination (Bähler et al., 1998).

JM8699 (*Pnmt81-cdc13+-internal sfGFPcp::his+)* was created by integrating plasmid pJM1794 (*pHis5Stu1-P81nmt1-Cdc13-sfGFPint*), which was generated by two steps of Gibson assembly. First, *Pcdc13-cdc13+-internal-sfGFPcp* was PCR amplified from genomic DNA from JM7788. The backbone fragment was amplified from pJM1492 (*pHis5Stu1*, Vjestica et al., 2020). Gibson assembly from these fragments was used to generate plasmid pJM1653 (*pHis5Stu1-Pcdc13-cdc13-sfGFPint*). Second, the *Pnmt81* promoter was PCR amplified from plasmid pJM240 (*pFA6a-kanMX6-p81nmt1-mCFP*). The backbone fragment, including *cdc13-*sfGFPint but excluding the endogenous *Pcdc13* promoter, was amplified from pJM1653. These products were combined by Gibson assembly to generate pJM1794, which was integrated into the *his5* locus of JM6437 (*his5-D21).* Correct colonies were those that grew on EMM-His medium. The resulting colonies were additionally screened by microscopy, where addition of thiamine repressed fluorescent Cdc13 expression.

### Microscopy

For all experiments fission yeast cells were first grown at 25°C in yeast extract supplemented with adenine, histidine, leucine, and uridine (YE4S).

*For glucose limitation experiments* (Fig. 2, 3; Fig.S1): After 8-16 hours of culturing in YE4S, cells were maintained in YE4S with 3% glucose (167 mM) or washed and moved into YE4S with 0.08% glucose (4.4 mM) or YE4S with 0.1% glucose (5.5 mM) and grown at 25°C for 36-48 hours to logarithmic phase prior to imaging.

*For nitrogen limitation experiments* (Fig. 4; Fig. S2): After 8-16 hours of culturing in YE4S, cells were washed and moved into Edinburgh Minimal Medium supplemented with adenine, histidine, leucine, and uridine (EMM4S) and grown at 25°C. After 8-16 hours of culturing in minimal media, cells were washed and moved into EMM supplemented with either 20 mM glutamate (EMM + glutamate), 20 mM proline (EMM + proline), or 20 mM isoleucine (EMM + isoleucine) and grown at 25°C for 36 hours to logarithmic phase prior to imaging.

All imaging was performed using a Nikon EclipseTi-2E microscope with a Yokogawa CSU-WI (Nikon Software) spinning disk confocal system. This system was equipped with a Photometrics PrimeBSI sCMOS camera (Eclipse Ti2; Nikon) and a 60× 1.4-NA CFI60 Apochromat Lambda S oil objective lens (Nikon). 405-, 488-, and 561-nm laser lines were utilized. smFISH imaging was performed with a 100x 1.4-NA CF160 Apochromat Lamda S oil objective lens (Nikon).

### Cell population studies

Doubling time of cell populations were determined from optical density (OD_600_) readings taken throughout an 8-hour time period at 25°C.

For static imaging experiments, cells were placed in a glass-bottom dish (#1.5, 35mm; ibidi) and covered with a YE4S (3% or 0.08% glucose) agar pad prewarmed to 25°C. Multiple fields of view per cell type were imaged within 2 hours at room temperature. Images were captured with 27 z-stacks and 0.2-μm step size using a spinning disk confocal microscope (described above) (Fig. 2; Fig. S1). For imaging Leptomycin B treated cells, 50 ng/µL LeptB was added to cells 30 or 60 mins prior to mounting on a glass-bottom dish and incubated shaking at 25°C (Figure 1H). All experiments were repeated three times; representative graphs are from one biological replicate.

### Single-cell studies

The ONIX microfluidics perfusion system from CellASIC (Millipore, CellASIC ONIX, 3.5 – 5.5 μm Y04C-02) was used for single cell imaging of Cdc13 accumulation over time. Prior to imaging, Y04C microfluidics plates were set up as described in the Y04C-02-5PK User Guide. 250 µL of cell culture was loaded at a density of approximately 1.2 x 10^6^ cells/ mL into the cell inlet. Depending on the experiment, the relevant media was loaded into all solution inlets, and a flow rate of 2 psi (13.8 kPa) was maintained throughout the experiment. All cells were imaged in the 3.5 µm chamber. Timelapse imaging began 1 hour after loading the microfluidics plate and continued for 12-24 hours at room temperature (∼23-25°C). The duration of each solution inlet flow was determined based on the number of consecutive cell cycles desired. Images were acquired with 11 z-stacks and 0.5 µm steps every 10 mins using spinning disk confocal microscopy. Glucose and nitrogen limitation experiments were repeated three times; representative graphs are from one biological replicate (Fig. 2, 3H-I, 4; Fig. S1, S2).

*For cycloheximide experiments* (Fig. 1C-E, Fig. S1C): The microfluidics plates were set up as described above. The first two solution inlets only contained media while the remaining 4 solution inlets were contained media with EtOH (vehicle control) or 1000 µg/mL cycloheximide (CHX; originally resuspended in EtOH). Timelapse imaging proceeded as described above. Nuclear measurements of the 488 channel of wildtype cells exposed to 1000 µg/mL CHX over time was used for background subtraction to quantify Cdc13 half-life. We note that previous studies examining bulk cell populations estimated a short 30-90 minute half-life for Cdc13 (Esposito et al., 2022). However, we observed some cells treated with CHX still entered mitosis, when Cdc13 is rapidly degraded by Anaphase Promoting Complex (APC)-mediated proteolysis. Since time-dependent Cdc13 accumulation occurs during interphase, we omitted these cells that entered mitosis after cycloheximide addition.

*For thiamine repressible promoter experiments* (Fig. 1F-G, Fig. S1D): Agar pads were prepped as described in “Cell populations studies.” Cell cultures were started in YE4S at 25°C and moved to EMM4S to grow for at least 20 hours without the presence of thiamine. Prior to imaging, cells were resuspended in EMM4S + 2 µg/mL thiamine, plated to a 35mm glass-bottom dish, and covered with a prewarmed agar pad. Images were acquired 5 mins after thiamine addition, with 11 z-stacks and 0.5 µm steps every 10 mins for 5 hours at room temperature using a spinning disk confocal microscope (described above). These cells expressed two copies of Cdc13: the first copy expressed by the P81nmt1 promoter, and the second copy expressed by the endogenous promoter. Therefore, repression of P81nmt1-driven expression did not prevent entry into mitosis. We only analyzed cells prior mitotic entry due to rapid Cdc13 degradation during anaphase.

### Single-Molecule Fluorescence In-Situ Hybridization (smFISH)

Quantification of mRNA by smFISH was performed as previously described (Esposito et al., 2022), with minor modifications. Briefly, approximately 2 x 10^8^ cells were fixed with 4% paraformaldehyde at room temperature for 30 minutes. Cells were then washed three times with ice-cold Buffer B (1.2 M sorbitol, 100 mM potassium phosphate buffer pH 7.5) and stored at 4°C overnight. To digest cell walls, samples were resuspended in spheroplast buffer (1.2 M sorbitol, 0.1 M postassium phosphate, 20 mM vanadyl ribonuclease complex (NEB S1402S), 20 μM beta-mercaptoethanol) containing 0.002% 100 T zymolyase (US Biological, Z1005) for approximately 1 hour. Following three washes in Buffer B, cells were gently resuspended and incubated for 20 minutes in 1 mL of 0.01% Triton X-100 in 1x PBS. Cells were washed three times in Buffer B, followed by one wash in 10% formamide in 2x saline-sodium citrate (SSC) pH 7.

For each sample, approximately 20–25 ng of Stellaris CAL Fluor 610 probes targeting the mEGFP coding sequence (Table S2) was mixed with 2 µL of yeast tRNA (Thermo Fisher Scientific, AM7119), 2 µL of Herring Sperm DNA (Sigma, D7290), and 45 µL of Buffer F (20% formamide, 10 mM sodium-phosphate buffer pH 7.2). This mixture was heated at 95°C for 3 minutes and allowed to cool to room temperature before combining with 50 µL of Buffer H (4x saline-sodium citrate (SSC) buffer, 4 mg/ml acetylated BSA (Thermo Fisher Scientific, 2614G1), 20 mM vanadyl ribonuclease complex) to create the final probe solution. Cells were resuspended in 100 µL of probe solution and incubated at 37°C overnight wrapped in aluminum foil.

Cells were washed with 10% formamide/2x SSC followed by 0.1% Triton X-100/2x SSC. For nuclear staining, cells were incubated in the dark in 1x PBS with 0.5 μg/ml Hoechst 33342 (Cell Signaling, 4082S) for 10 minutes and then washed with 1x PBS. Cell pellets were gently resuspended in SlowFade Diamond Antifade Mountant (Thermo Fisher Scientific, S36972) prior to mounting on glass slides using #1.5 glass coverslips. For each channel, images were acquired every 0.2 μm for a total of 6 μm in Z. Images were deconvolved and analyzed using FISHquant software (Mueller et al., 2013) following the suggestions outlined in the FISHquant documentation. Successful RNA spot detection in each cell was verified by manual inspection.

### Cell size homeostasis studies

Cells were imaged in microfluidic flow chambers (Millipore, CellASIC ONIX, 3.5 – 5.5 μm Y04C-02) as described above. Brightfield images of cells were acquired with 11 z-stacks and 0.5 µm steps every minute using a spinning disk confocal microscope (described above) (Fig. 3I). Cell size at birth and division and cycle time were analyzed using Cell ACDC (Padovani et al., 2022). Cells were segmented from brightfield images signal using the YeaZ-v2 model (Dietler et al., 2020) with default parameters, including a threshold value of 0.4 and a minimum distance of 10 pixels. All automatic segmentations and cell labels were then manually reviewed, corrected, and verified using the Cell-ACDC graphical user interface.

### Image Analysis

All image analysis was performed using ImageJ unless otherwise specified. Static and timelapse imaging was analyzed for geometry measurements and fluorescent intensity as described in “Cell and nuclei segmentation”, “Cell geometry measurements”, “Nuclear size measurements”, and “Measurement of fluorescent intensity” from (Miller et al., 2023). Measurement of surface area is calculated using the equation of a cylinder with hemispherical ends (Miller et al., 2023; Pan et al., 2014; Facchetti et al., 2019).

Background subtraction for timelapse fluorescence intensity measurements was completed by subtracting the average nuclear or whole cell GFP fluorescence of untagged wild type cells, unless otherwise specified.

### Immunoblotting

Fission yeast whole cell lysates were prepared by growing cells in their respective media to mid-logarithmic phase. Two OD of cells (typically 10 mL of cells at OD_600_ of 0.2) were harvested and snap-frozen in liquid nitrogen. Cell pellets were resuspended in SDS-PAGE sample buffer (15% glycerol, 4.5% SDS, 97.5 mM Tris pH 6.8, 10% 2-mercaptoethanol, 50 mM β-glycerophosphate, 50 mM sodium fluoride, 5 mM sodium orthovanadate, 1x EDTA-free protease inhibitor cocktail (Sigma Aldrich)). The samples were lysed with acid-washed glass beads (Sigma Aldrich) in a MiniBeadBeater-16 (BioSpec, Bartlesville, OK) for 2 minutes at 4°C. Cell lysates were then incubated at 99°C for 5 minutes. Cell lysates were briefly centrifuged, and the supernatant was isolated as the clarified lysate. 5 µL of the cell lysates were separated by SDS-PAGE and transferred to a nitrocellulose membrane using the Trans-blot Turbo Transfer System (Bio-Rad). The antibodies listed below were used for the detection of proteins by western blotting. Blots were washed multiple times in 1X TBST and once in 1X TBS and signal was detected using a LICOR Odyssey CLx.

*For cycloheximide protein degradation experiments* (Fig. S1B-C): The temperature sensitive mutant strain *cdc25-22* was cultured at the permissive temperature (25°C) for 24 hours before shifting to the restrictive temperature (36.5°C) for 3 hours. Successful and complete G2 arrest was confirmed by septation index prior to cycloheximide addition, and at every experimental timepoint. 1000 µg/mL cycloheximide was added. Two OD_600_ samples were harvested and snap-frozen in liquid nitrogen prior to treatment and at the following timepoints post cycloheximide addition: 15, 30, 60, 90, 180, 360 mins. Experiment was repeated three times.

To quantify Cdc13 levels, a segmented mask of the bands was created. The relative band intensities were determined following background subtraction and normalization to total protein levels using Ponceau stain. Error bars indicate standard deviation.

*For Cdk1 phosphorylation experiments* (Fig. 3): Cells were cultured, and lysates were prepared as described above. Experiments were repeated three times. To quantify Cdk1 phosphorylation levels, segmented masks of the bands on were created. The relative band intensities of Cdk1 phospho-sites were determined following background subtraction and normalization to total Cdk1 protein levels. Error bars indicate standard deviation.

*Primary antibodies were used at the following concentrations:* anti-Cdc13, 1:1000 [Cdc13 (6F11/2), mouse monoclonal; NB200-576, Novus Biologicals]; anti-Cdk1-Y15P, 1:3000 [phospho-Cdc2 (Y15), rabbit polyclonal; #9111, Cell Signaling Technology]; anti-Cdk1, 1:10000 [Cdc2 p34 (Y100.4), mouse monoclonal; sc-53217, Santa Cruz Biotech]; anti-Cdk1-T161P for the detection of *S. pombe* Cdk1-T167, 1:1000 [Phospho-cdc2 (Thr161), rabbit polyclonal; #9114, Cell Signaling Technology].

*Secondary antibodies were used at the following concentrations*: goat anti-mouse secondary antibody, 1:6600 (IRDye 800CW Goat anti-Mouse IgG, 926-32210, LICOR); goat anti-rabbit secondary antibody, 1:10000 (IRDye 800CW Goat anti-Rabbit IgG, 926-32211, LICOR).

### Statistical Analysis

All statistics and graphs were generated using Prism10 (GraphPad software). Unpaired t-tests were performed to determine statistical difference between two sets of data. p-values < 0.05 were considered statistically significant. Non-linear regression (one phase decay) analysis was used to estimate the Cdc13 half-life in high and low glucose concentrations from cycloheximide experiments. Linear regression analysis was used to estimate Cdc13 half-life in high glucose from Pnmt81 promoter shut-off experiment and to obtain slope values for size homeostasis plots.

### Outlier Analysis

An outlier analysis was performed on Cdc13 nuclear export data (Fig. 1H) using Prism10’s “Identify outliers” function. The ROUT method was utilized with a False Discovery Rate (Q) set to 10%. One outlier was identified and removed in the 9-12 µm size bin of the methanol control group, and two outliers were identified and removed in the 9-12 µm size bin of the Leptomycin B treatment group.

### Binning Analysis

For *cdc13* mRNA FISH experiments cells were binned into 1 µm cell length bins (Fig. 1B). For glucose limitation experiments cells were binned by their surface area with the same number of cells in each bin using the quantile function included in R (Fig. S1G-H).

### Mathematical Modeling Mechanism of cell size regulation

The model described here links cell growth to the dynamics of nuclear Cdc13 and to size-dependent activation of Cdk1, providing a quantitative framework for how division timing is controlled. For simplicity, the cell cycle is represented by two functional states: G2 and M/G1/S. In the G2 phase, the cell elongates and accumulates nuclear Cdc13. During M/G1/S phase, growth is negligible, the nucleus divides, and cytokinesis occurs. Cell size at time *t* is denoted by *s*(*t*), and its dynamics is given by:

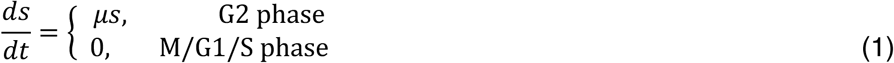

in which *μ* is the exponential growth rate. For simplicity, the explicit time dependence of all dynamical variables is omitted unless needed for clarity.

The nuclear concentration of Cdc13 is denoted by *x*(*t*). Cdc13 is imported to the nucleus at a constant rate throughout the cell cycle. During G2, its concentration is reduced by dilution due to cell growth. During M/G1/S, growth is negligible and the dilution term is absent. Thus,

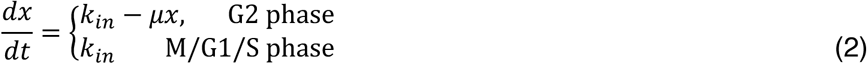

where *k*_in_ is the rate of nuclear Cdc13 accumulation. At entry into M/G1/S, nuclear Cdc13 is rapidly degraded and reset to a basal concentration *x_b_*, which is assumed to follow a gamma distribution with mean ⟨*x_b_*⟩ and squared coefficient of variation CV^2^ . This reset represents APC/C-mediated degradation. Note that throughout the paper ⟨ ⟩ is used to denote the mean value of a random variable.

Mitotic entry is controlled by the concentration of active Cdk1, denoted by *c*^∗^. In this model, active Cdk1 depends on nuclear Cdc13 concentration and on a size-dependent regulatory term representing the combined effect of Cdc25 and Wee1. The contribution of the available Cdk1 pool is incorporated implicitly into the effective scaling. Assuming that the underlying binding and activation steps are fast compared with cell growth and cell-cycle progression, active Cdk1 is described by the following multiplicative interaction:

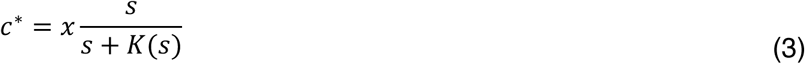

In this equation, size dependence of Cdk1 activation results from Cdc25 concentration increasing linearly with size and Wee1 activity decreasing with size as per a Hill function *K*(*s*) (Miller et al., 2023). During G2, equation 3 depends on *s*, while in M/G1/S it depends on *s*/2, since the cell contains two nuclei before cytokinesis and the relevant size-dependent regulation is assumed to act at the level of each daughter nucleus. The net effect of Wee1 activity is incorporated into the effective function:

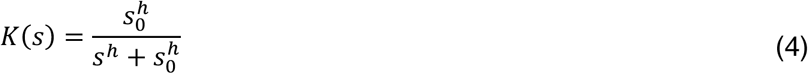

where *s*_0_ is the size saturation coefficient, and ℎ is the Hill coefficient. As cell size increases, *K*(*s*) decreases, increasing the fraction of active Cdk1.

Division is initiated when active Cdk1, shown in equation (3), reaches an activation threshold, denoted by *c*^†^. This threshold is assumed to be stochastic, drawn from a gamma distribution with mean B*c*^†^C and squared coefficient of variation CV^2^ . Crossing this threshold marks entry into M/G1/S, after which the following transitions occur. First, cell growth stops temporarily. Second, nuclear Cdc13 is rapidly degraded, so that *x* → *x_b_*, where *x_b_* is a small stochastic basal concentration. Third, the duration of M/G1/S phase is assumed to follow a gamma distribution with a given mean and variance. Finally, at cytokinesis, cell size is partitioned according to *s* → *βs*, where *β* is a stochastic partitioning factor drawn from a beta distribution with mean ⟨*β*⟩ and coefficient of variation CV^2^.

### Parameter fitting for cell size regulation under different glucose concentration

Parameter values were first determined for the high-glucose condition. Static single-cell measurements of nuclear Cdc13 concentration as a function of cell surface area (Fig. 2C) were used to constrain the Cdc13-related parameters. The parameters 〈*x_b_*〉 and CV^2^ were estimated from the minimum observed Cdc13 level, while *k_in_* was adjusted so that the model reproduced the observed range of Cdc13 concentration. The duration of M/G1/S was assumed to follow a gamma distribution with mean and variance chosen to approximately match the experimental measurements. The remaining parameters (*s*_0_, ℎ, 〈 *c*^†^〉 and CV^2^ ) for the high-glucose condition were estimated by minimizing the Maximum Mean Discrepancy (MMD) (Gretton et al., 2012) between the simulated and experimental joint distributions of surface area at birth and added surface area. The resulting fitted parameters are listed in Table 1 together with the parameters constrained directly from experimental data.

After estimating the parameters for the high-glucose condition, the low-glucose condition was considered. Experimental results indicated that the growth rate is reduced in low glucose. Accordingly, for the low-glucose condition, the growth rate was fixed from experimental data, and the activation threshold was re-estimated using the same MMD-based procedure, while the remaining parameters were kept at their high-glucose values (Table 1). Consistent with the experimental data, the simulations predicted size homeostasis in both high- and low-glucose conditions, as reflected by a negative slope when plotting surface area at birth and added surface area. The model also predicted higher Cdc13 concentration in low-glucose cells than in high-glucose cells, consistent with the experimental trend (Fig. 2, 3). Finally, the model predicted a reduction in cell size in low glucose conditions, consistent with experiments (Fig. 2, 3).

## Supplemental Material

Fig. S1 shows Cdc13 accumulation in the nucleus remains time dependent under glucose limitation, and *cdc13+* RNA concentration measurements. Fig. S2 shows that Cdc13 accumulation is not time-dependent under nitrogen limitation. Table S1 describes yeast strains and smFISH primers used in this study.

## Data Availability

All primary data are available upon request.

## Acknowledgements

We thank members of the Moseley lab for helpful comments on the manuscript; as well as the Biomolecular Targeting Core (BioMT) (P20-GM113132) for use of equipment. We thank Kristi Miller for support and ideas in the early stages of this project. This work was supported by grants from the National Institutes of General Medical Sciences (NIGMS) (R35GM149248) to J.B.M and (1F31GM156081) to S.E.V.

## Supplemental Figure Legends

**Figure S1.** (A) Total nuclear Cdc13-sfGFP (au) of single cells treated with cycloheximide (1000 µg/mL) over time. Data were obtained from the same cells analyzed in Fig. 1E. 0 mins indicates cycloheximide addition. Timepoints were fit to a one-phase decay function to determine half-life of Cdc13 (t_1/2_ = 169.5 mins). (B) Immunoblot against Cdc13 of whole cell extracts from *cdc25-22* cells grown in 3% glucose. Cells were shifted to 36.5°C for 3 hours to arrest in G2 phase, and then treated with cycloheximide (1000 µg/mL). Samples were acquired at the indicated timepoints. (C) Quantification of Cdc13 degradation in G2-arrested cells normalized against total protein levels. Timepoints were fit to a linear function to determine half-life of Cdc13 (t_1/2_ = 116 mins). (D) Total nuclear Cdc13-sfGFP (au) of *Pnmt81-cdc13-sfGFP* single cells over time following addition of thiamine (2 µg/mL) (n = 20 cells). Data were obtained from the same cells analyzed in Fig. 1G. 0 mins indicates thiamine addition. (E) Doubling time (hrs.) of cell population grown in 3% glucose media or 0.08% glucose media at 25°C. (F) Cdc13-sfGFP whole cell concentration in cells grown in 3% and 0.08% glucose plotted by cell size (µm^2^). Same cells that are plotted in Fig. 1C. (G) Cdc13-sfGFP nuclear concentration for cells grown in media with 3% and 0.08% glucose binned by cell size. Same cells that are plotted in Fig. 1C and S1F. (H) Cdc13 concentration in the whole cell for WT cells grown in media with 3% and 0.08% glucose binned by cell size. Same cells that are plotted in Fig. 1C and S1F. (I) Total nuclear Cdc13-sfGFP of single cells from timelapse imaging for cells grown in 3% or 0.08% glucose plotted against time since birth. Same cells that are plotted in Fig. 2F-I. Error bars indicate standard deviation. (J - K) Cdc13-sfGFP concentration in the nucleus of single cells from timelapse imaging for cells grown in 3%, 0.1%, or 0.08% glucose. Concentrations are plotted against either cell size (µm^2^) (panel J) or time since birth (panel K). Error bars indicate standard deviation. (n = 10 cells per condition). (L-M) Averaged Cdc13-sfGFP mean nuclear concentration (au) from timelapse imaging of WT and *cdr2Δ* cells grown in media with 0.08% glucose plotted either by cell size (panel L) or time since birth (panel M) (n = 10 cells per condition). The same cells are plotted in panels L and M. Error bars indicate standard deviation. (N) Representative brightfield and Cdc13-sfGFP fluorescence images of an interphase WT cell grown in a microfluidics device with constant access to rich media with 0.08% glucose over time after treatment with cycloheximide. (O) Nuclear Cdc13-sfGFP concentration (au) of single cells treated with cycloheximide (1000µg/mL) over time (0.08% glucose n = 18). 0 mins indicates cycloheximide addition. Timepoints were fit to a one-phase decay function to determine half-life of Cdc13 in cells grown in media with 0.08% glucose (t_1/2_ = 165.1 mins). (P) Representative brightfield and smFISH *cdc13-mEGFP* transcript fluorescence images of *cdc13-mEGFP-3’UTR (cdc13)* cells grown in 0.08% glucose. Hoechst stain marks the nucleus (scale bar = 10 µm). (Q*) cdc13+* mRNA concentration (# molecules per cell length) for cells grown in 3% or 0.08% glucose (3% glucose n = 135 cells, 0.08% glucose n = 164 cells). (R) *tps2+* mRNA concentration (# molecules per cell length) for cells grown in 3% or 0.08% glucose (3% glucose n = 112 cells, 0.08% glucose n = 84 cells). (S) *leu2+* mRNA concentration (# molecules per cell length) for cells grown in 3% or 0.08% glucose (3% glucose n = 126 cells, 0.08% glucose n = 126 cells). (T) *his3+* mRNA concentration (# molecules per cell length) for cells grown in 3% or 0.08% glucose (3% glucose n = 115 cells, 0.08% glucose n = 77 cells). ** p < 0.01; **** p < 0.0001 by unpaired t-test.

**Figure S2. Analysis of Cdc13 accumulation for cells grown in different nitrogen sources.** (A) Doubling time (hrs.) of cell population grown in media supplemented with either glutamate or proline at 25°C. (B) Cell size at division (µm^2^) of *cdc13-sfGFP* cells grown in minimal media supplemented with glutamate or proline. Same cells that are plotted in Fig. 4C. (C) Total nuclear Cdc13-sfGFP of single cells from timelapse imaging for cells grown in minimal media supplemented with glutamate or proline plotted against time since birth. Same cells that are plotted in Fig. 4C-F. Error bars indicate standard deviation. (D) Averaged Cdc13-sfGFP concentration (au) in the nucleus of single cells grown in minimal media supplemented with glutamate, isoleucine, or proline. Concentrations are plotted against cell size (µm^2^). Error bars indicate standard deviation. (E-F) Averaged Cdc13-sfGFP mean nuclear concentration (au) from timelapse imaging of WT and *cdr2Δ* cells grown in minimal media supplemented with proline either by cell size (panel E) or time since birth (panel F). Same cells are plotted in Fig. S2E and F. Error bars indicate standard deviation. *** p <0.001 by unpaired t-test.

