## Supplementary figures and images for "The role of cell growth rate on accumulation of the mitotic cyclin Cdc13 in fission yeast"

### Supplemental Figure S1

#### Supplemental Figure 1.

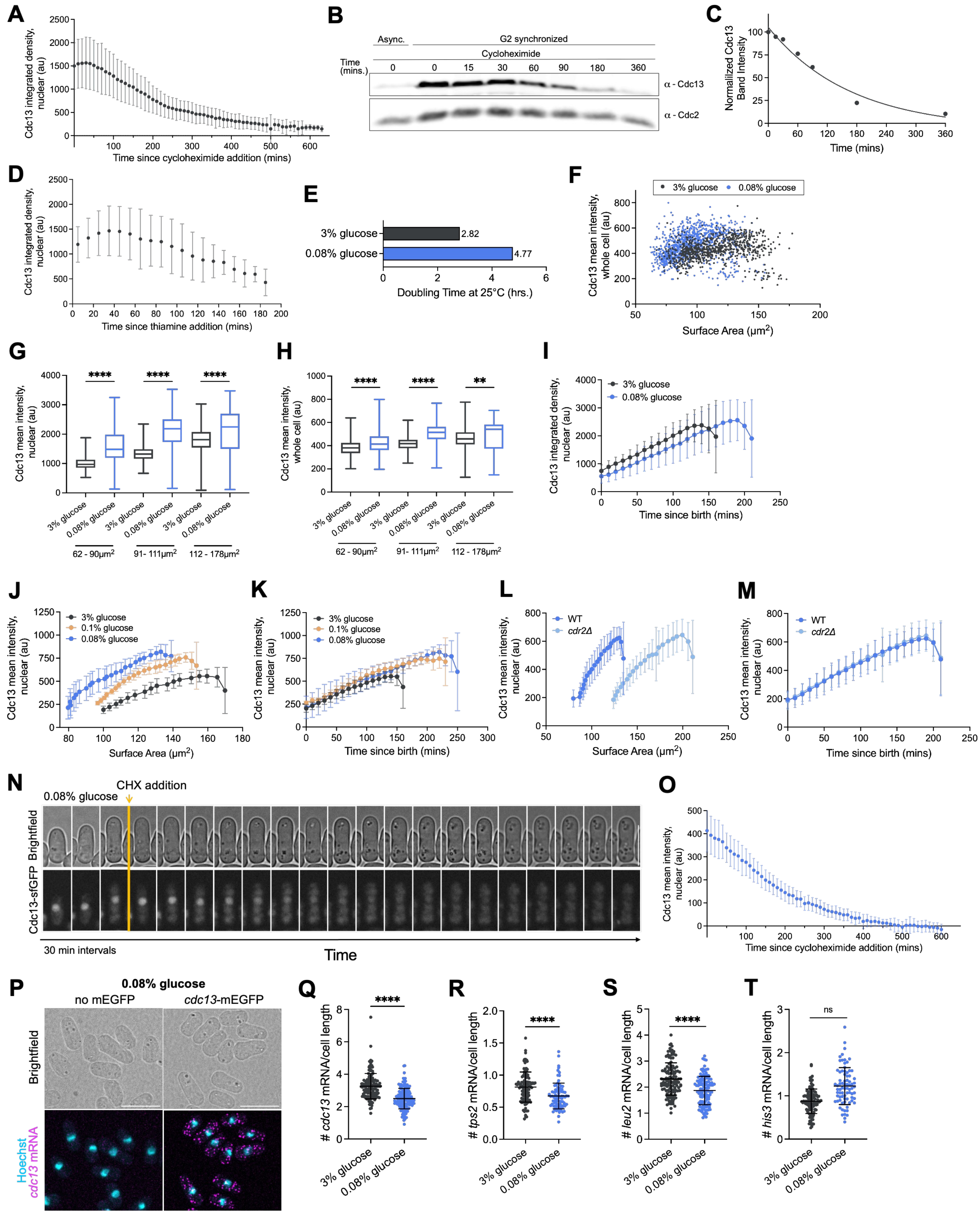

### Supplemental Figure S2

#### Supplemental Figure 2.

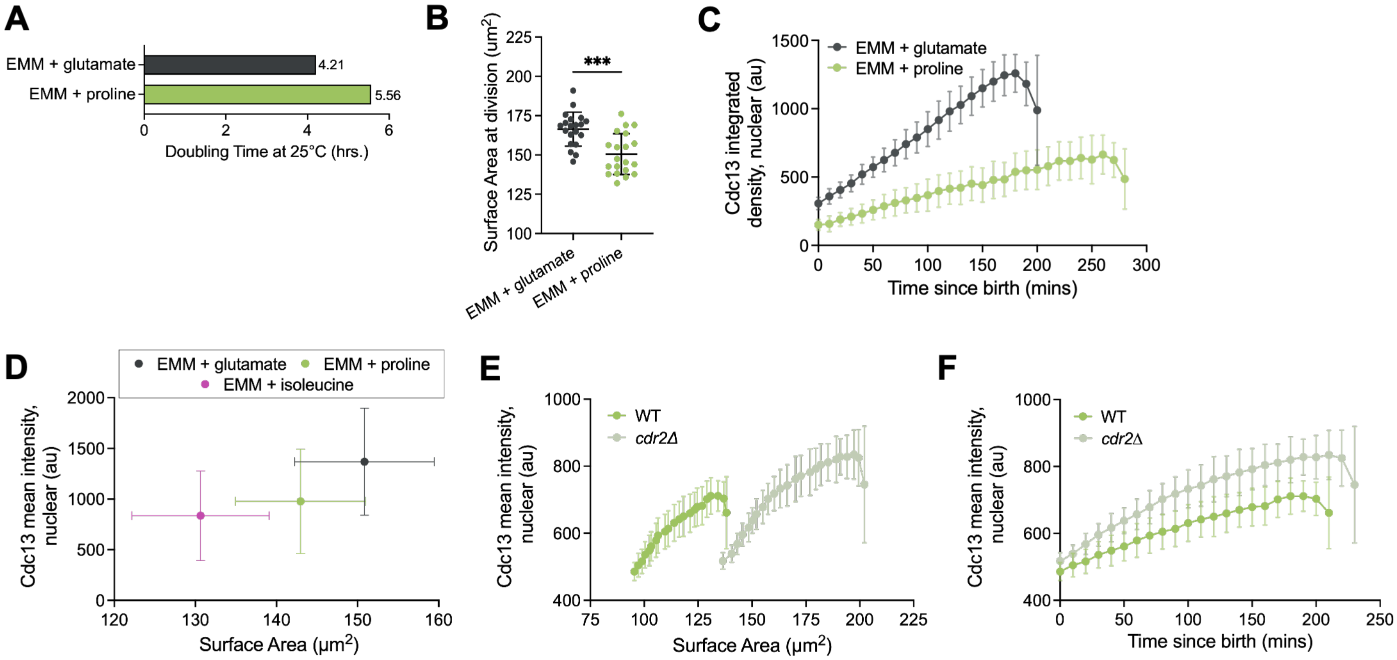
