## Supplemental Table S1 for "The role of cell growth rate on accumulation of the mitotic cyclin Cdc13 in fission yeast"

**Table S1. Yeast Strains**

| Strain | Description | Source |
|---|---|---|
| JM14 | cdc25-22 h+ | Lab stock |
| JM366 | 972 h- | Lab stock |
| JM367 | 975 h+ | Lab stock |
| JM6437 | his5-D21 h- | Lab stock |
| JM6907 | cdc2-mNeonGreen::hphR lys3+:$P_{tdh1}$*-NLS-linker-mTagBFP2-$T_{tdh1}$::kanMX h- | Lab stock |
| JM7748 | lys3+:$P_{tdh1}$*-NLS-linker-mTagBFP2-$T_{tdh1}$:kanMX h- | Lab stock |
| JM7788 | cdc13+-internal-sfGFPcp lys3+:$P_{tdh1}$*-NLS-linker-mTagBFP2-$T_{tdh1}$:kanMX h+ | This study |
| JM8194 | cdr2Δ::natMx cdc13+-internal-sfGFPcp lys3+:$P_{tdh1}$*-NLS-linker-mTagBFP2-$T_{tdh1}$:kanMX h? | This study |
| JM8496 | cdc13-mEGFP-3'UTR(cdc13)::kanMX h- | This study |
| JM8699 | $P_{nmt81}$-cdc13+-internal-sfGFPcp::his+ h? | This study |
| JM8981 | his3-mEGFP::kanMx h- | This study |
| JM8982 | tps2-mEGFP::kanMx h- | This study |
| JM8995 | leu2-mEGFP::kanMx h- | This study |

**Table S2. smFISH probes targeting the mEGFP coding sequence**

| Probe # | |
|---|---|
| 1 | gtgaaaagttcttctccttt |
| 2 | acaagaattgggacaactcc |
| 3 | ccattaacgtcaccatctaa |
| 4 | ccattaacgtcaccatctaa |
| 5 | gtaagttttccgtatgttgc |
| 6 | gtagttttccagtagtgcaa |
| 7 | acaagtgttggccatggaac |
| 8 | acaccataagtcagagtagt |
| 9 | ctgggtatcttgaaaagcat |
| 10 | agtcatgctgtttcatatga |
| 11 | cgggcatggcactcttgaaa |
| 12 | ttctttcctgtacataacct |
| 13 | gttcccgtcatctttgaaaa |
| 14 | tgacttcagcacgtgtcttg |
| 15 | taacaagggtatcaccttca |
| 16 | ataccttttaactcgattct |
| 17 | agaatgtttccatcttcttt |
| 18 | ttgtattccaatttgtgtcc |
| 19 | gtctgccatgatgtatacat |
| 20 | ctttgattccattcttttgt |
| 21 | tccatcttcaatgttgtgtc |
| 22 | gttgataatggtctgctagt |
| 23 | catcgccaattggagtattt |
| 24 | tgtctggtgaaaggacaggg |
| 25 | cttagattgtgtggacaggt |
| 26 | tcttttcgttgggatctttc |
| 27 | tcaagaaggaccatgtggtc |
| 28 | atcccagcagctgttacaaa |
| 29 | tatagttcatccatgccatg |
